# Difluoromethylornithine rebalances aberrant polyamine ratios in Snyder-Robinson syndrome: mechanism of action and therapeutic potential

**DOI:** 10.1101/2023.03.30.534977

**Authors:** Tracy Murray Stewart, Jackson R. Foley, Cassandra E. Holbert, Maxim Khomutov, Noushin Rastkari, Xianzun Tao, Alex R. Khomutov, R. Grace Zhai, Robert A. Casero

## Abstract

Snyder-Robinson Syndrome (SRS) is caused by mutations in the spermine synthase (*SMS*) gene, the enzyme product of which converts the polyamine spermidine into spermine. Affecting primarily males, common manifestations of SRS include intellectual disability, osteoporosis, hypotonic musculature, and seizures, along with other, more variable symptoms. Currently, medical management focuses on treating symptoms without addressing the underlying molecular cause of the disease.

Reduced SMS catalytic activity in cells of SRS patients causes the accumulation of spermidine, while spermine levels are reduced. The resulting exaggeration in spermidine-to-spermine ratio is a biochemical hallmark of SRS that tends to correlate with symptom severity in the patient. Our studies aim to pharmacologically manipulate polyamine metabolism to correct this polyamine imbalance and investigate the potential of this approach as a therapeutic strategy for affected individuals.

Here we report the use of 2-difluoromethylornithine (DFMO; eflornithine), an FDA-approved inhibitor of polyamine biosynthesis, in re-establishing normal spermidine-to-spermine ratios in SRS patient cells. Through mechanistic studies, we demonstrate that, while reducing spermidine biosynthesis, DFMO also stimulates the conversion of existing spermidine into spermine in cell lines with hypomorphic variants of *SMS*. Further, DFMO treatment induces a compensatory uptake of exogenous polyamines, including spermine and spermine mimetics, cooperatively reducing spermidine and increasing spermine levels. In a *Drosophila* SRS model characterized by reduced lifespan, adding DFMO to the feed extends lifespan. As nearly all known SRS patient mutations are hypomorphic, these studies form a foundation for future translational studies with significant therapeutic potential.

## Introduction

Spermine synthase (SMS) (EC 2.5.1.22) is the sole source of *de novo* spermine (SPM) generation in mammalian cells (1). The final biosynthetic step of the mammalian polyamine pathway, SMS catalyzes the transfer of an aminopropyl group from decarboxylated S-adenosylmethionine (dcSAM) to spermidine (SPD), thereby forming SPM (**Fig. 1**) (2). Germline mutations of the *SMS* gene result in Snyder-Robinson syndrome (SRS; OMIM #309583), a recessive X-linked intellectual disability syndrome predominantly affecting males (3-5) and associated with developmental delay, osteoporosis, muscle hypotonia, seizures, and a variety of less common symptoms (6,7). Medical management of SRS targets these manifestations (8), as there is no current treatment that targets the underlying biochemical cause of the disease.

**Fig. 1.**
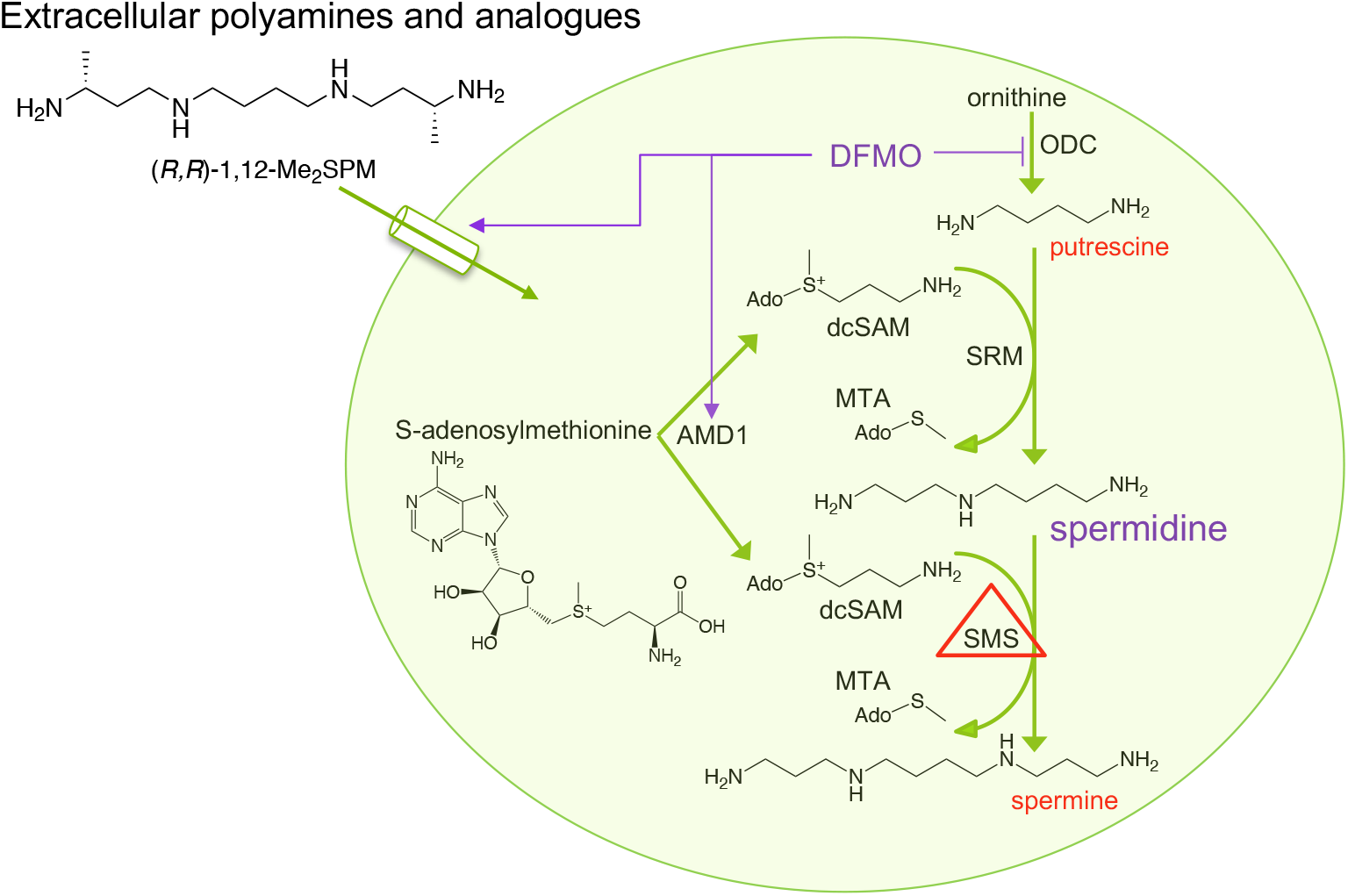
Effects of hypomorphic SMS and DFMO on the mammalian polyamine metabolic pathway in SRS patient-derived cells. Putrescine is derived from ornithine via ornithine decarboxylase (ODC), the first rate-limiting step in polyamine biosynthesis. S-adenosylmethionine decarboxylase (AMD1) activity provides the aminopropyl donor dcSAM for use by spermidine synthase (SRM) and spermine synthase (SMS) to create spermidine and spermine, respectively. In SRS, hypomorphic SMS function causes spermidine accumulation while spermine levels are reduced. DFMO is an irreversible suicide inhibitor of ODC. This block of polyamine biosynthesis gradually reduces polyamine pools, triggering compensatory uptake of extracellular polyamines and induction of AMD1 activity that favors conversion into the higher polyamines spermidine and spermine. MTA, methylthioadenosine.

Despite extracellular sources of SPM, including dietary consumption (9), a defining characteristic of SRS-affected cells and tissues is an elevated SPD/SPM ratio that does not appear to benefit from dietary SPM supplementation (5,6,10-16). The degree to which this ratio is altered in individual patients depends on the specific *SMS* mutation and appears to correlate with SMS enzymatic activity as well as phenotype severity, though existing data are insufficient to conclude a genotype-phenotype correlation (14).

The mammalian polyamines, including SPM, SPD, and their precursor putrescine (PUT), contribute to critical cellular functions that necessitate strict homeostatic control over their intracellular levels (1). Polyamine transport is generally downregulated in the presence of high intracellular polyamine concentrations (17), suggesting that SRS-affected cells might not transport extracellular SPM. However, we have shown that SPM transport activity in SRS patient-derived lymphoblastoid and fibroblast cells is as functional as that of wild-type *SMS* cells (18,19). Importantly, in response to SPM import, SRS-affected cells reduce their excessive SPD pools to within normal levels, indicating that polyamine homeostatic control mechanisms also remain functional. However, the fact remains that SPM is a common dietary component that fails to rescue the SRS phenotype *in vivo*.

We therefore proposed the use of 2-difluoromethylornithine (DFMO) as a treatment strategy targeting the aberrant SPD/SPM ratio by reducing spermidine biosynthesis while enhancing SPM production and extracellular SPM import. DFMO is a well-studied, clinically approved, irreversible inhibitor of ornithine decarboxylase (ODC), the first rate-limiting step of polyamine biosynthesis (**Fig. 1**)(20-22). ODC activity produces the diamine PUT, which is the precursor for SPD production via SPD synthase (SRM). Importantly for our studies, cells respond to this inhibition of polyamine biosynthesis by upregulating two compensatory mechanisms: 1) increased production of the aminopropyl donor, dcSAM, to facilitate synthesis of SPM, and 2) increased uptake of polyamines from the extracellular environment. Through these mechanisms, we reasoned that exposing SRS-affected cells to DFMO would improve the aberrantly elevated SPD/SPM ratio. We have therefore used lymphoblastoid and fibroblast cell lines from SRS patients as well as a *Drosophila* model of SRS to investigate the effects of DFMO exposure on proliferation, intracellular polyamine concentrations, extracellular polyamine uptake, polyamine metabolic enzyme activity, and phenotype reversal. Together, our results indicate that DFMO represents a potential treatment for SRS patients that targets the underlying biochemical aspects of the pathology. As DFMO is an FDA-approved drug with a history of safe administration, our results strongly support its further clinical development in the context of SRS.

## Results

### DFMO reduces the SPD/SPM ratio in SRS patient cells

As DFMO inhibits the first step in polyamine biosynthesis, we anticipated a reduction in intracellular SPD pools in SRS cells. Analyses of SRS patient-derived lymphoblastoid and fibroblast cell lines exposed to DFMO revealed dose-dependent reductions in SPD/SPM ratios that reached those determined in cells with wildtype *SMS* (**Fig. 2A** and **S1)**. As expected, decreased SPD concentrations contributed to this reduction. However, our data also indicated increases in SPM concentrations, suggesting enhanced conversion of SPD into SPM by these defective SMS enzymes in the presence of DFMO.

**Fig. 2.**
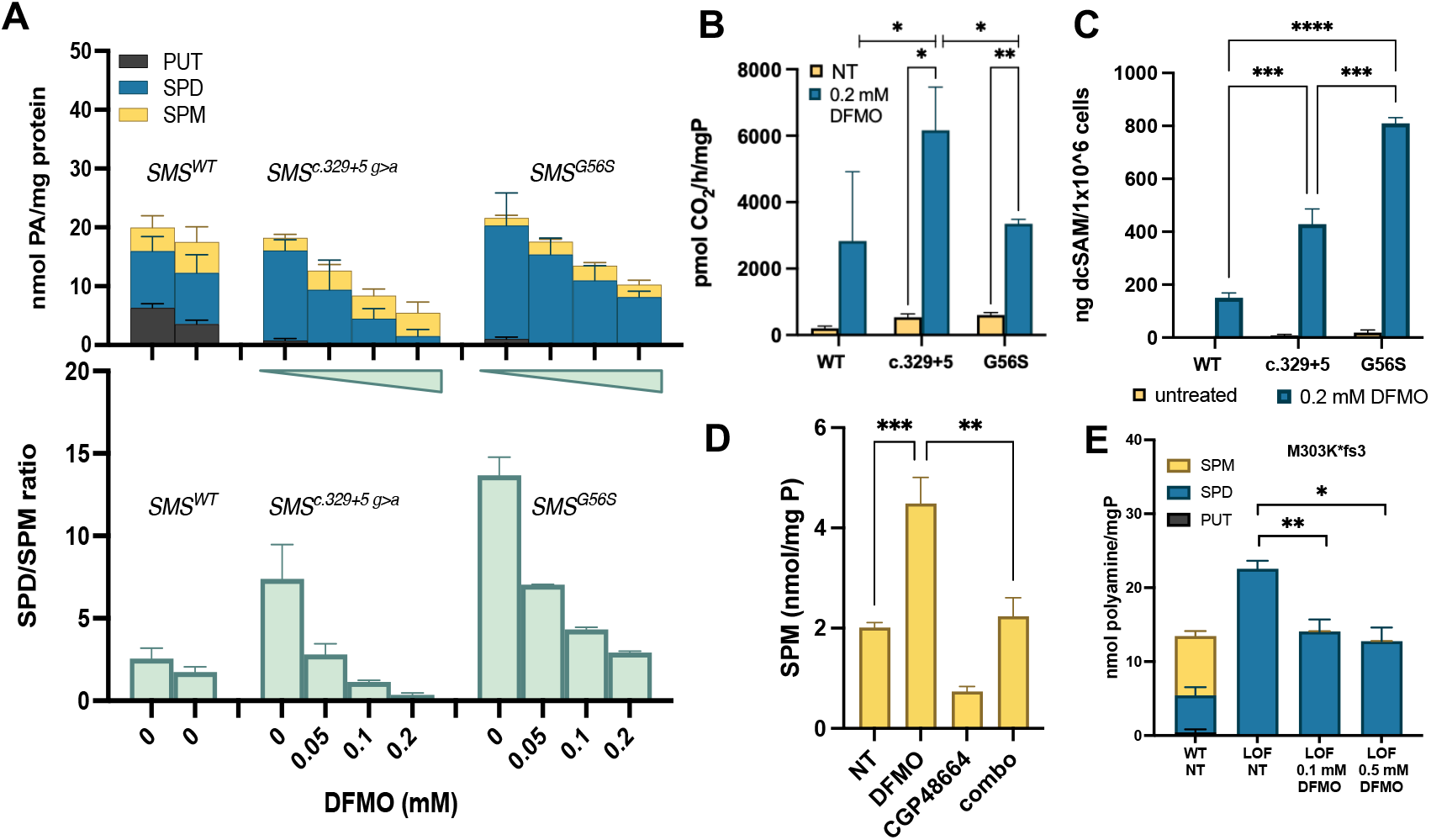
Mechanism of action of DFMO in SRS patient-derived lympoblastoid cells. DFMO treatment reduces intracellular SPD levels and increases SPM **(A top**), correcting the SPD/SPM ratio to *SMS*^*WT*^ levels (**A bottom**). DFMO treatment (96 h, 0.2 mM) induces AMD1 activity (**B**) and increases intracellular concentrations of dcSAM (**C**). Cotreatment with an AMD1 inhibitor (CGP48664) prevents the DFMO-mediated production of SPM in *SMS*^*G56S*^ cells (**D**), supporting increased availability of the aminopropyl donor (dcSAM) to the hypomorphic SMS enzyme as the mechanism of action. SRS patient cells with complete loss of SMS function (*SMS*^*M303K*fs3*^) fail to produce SPM in the presence of DFMO (**E**). All values were calculated from at least duplicate determination of at least 2 biological replicates.

In response to perturbations in polyamine biosynthesis, cells employ compensatory homeostatic control mechanisms (23). We considered that SRS-affected cells are likely responding to ODC inhibition by upregulating a second biosynthetic enzyme, S-adenosylmethionine decarboxylase (AMD1, **Fig. 1**). Treatment with DFMO (96 h, 0.2 mM) induced AMD1 activity in lymphoblastoid cells from SRS patients (**Fig. 2B)**, suggesting increased availability of the aminopropyl group donor, dcSAM, to the hypomorphic SMS enzymes. LC/MS analysis of intracellular dcSAM levels further support this hypothesis: dcSAM levels were elevated with DFMO treatment in all cell lines examined, with significantly greater levels evident in the SRS lines compared to WT (**Fig. 2C**). In the absence of DFMO, dcSAM was below the limit of detection in cells with WT *SMS*, while SRS patient lymphoblasts contained readily measurable amounts (**Fig. S2A**), consistent with a previous report (24). Levels of unmodified SAM did not differ among the untreated cell lines. However, treatment of the *SMS*^*G56S*^ SRS line with DFMO was associated with a reduction of SAM concentrations (**Fig. S2B**). Further, cotreatment with the AMD1 inhibitor CGP48664 (25) completely blocked the DFMO-mediated increase in SPM production in SRS cells (**Fig. 2D)**, supporting a critical role for AMD1 induction in the response to DFMO. To further confirm our hypothesis that DFMO facilitates the conversion of SPD to SPM in SRS cells containing hypomorphic variants of the SMS protein, we exposed SRS cells containing a complete loss-of-function *SMS* variant to DFMO (**Fig. 2E)**. Consistent with DFMO-mediated inhibition of biosynthesis, SPD was reduced in a dose-dependent manner; however, no SPM was evident in these cells regardless of DFMO dosage. Together, these data indicate a mechanism of action for DFMO in cells with hypomorphic SMS that includes reduced levels of SPD in conjunction with AMD1-mediated increased availability of dcSAM that facilitates SPM biosynthesis by the defective SMS enzyme (**Fig. 1**).

### SMS variants alter sensitivity to DFMO-mediated growth inhibition

DFMO can be anti-proliferative, particularly in rapidly dividing cells with increased dependence on polyamines. SRS or healthy donor fibroblast cell lines were incubated for 96 hours in the presence of increasing concentrations of DFMO (**Fig. 3A)**. Cell lines chosen for these studies represented severely affected (I150T, P112L) and more mildly affected (c.329 + 5g>a) patients with varying elevations in ratios resulting from the individual *SMS* mutations (19). Despite similar growth rates, these hypomorphic SRS cell lines were found to be significantly more resistant to the growth inhibitory effects of DFMO than WT cells from healthy donors (shown are combined data from 2 different WT cell lines). In contrast, cells with a complete loss-of-function (LOF; M303K*fs3) mutation in *SMS* demonstrated increased sensitivity to DFMO compared to WT cells. Specifically, the hypomorphic cell lines differed in the maximum inhibition of growth caused by DFMO exposure, ranging from approximately 10 to 27%, while WT cells demonstrated a maximum of approximately 46%. Although the maximum growth inhibition of the LOF mutant was similar to that of WT (∼50%), a left-shifted curve was observed that was reflected in a reduction of the IC_50_ value from 79 μM in *SMS*^WT^ cells to 27 μM in the LOF mutant line (**Table 1**).

**Fig. 3.**
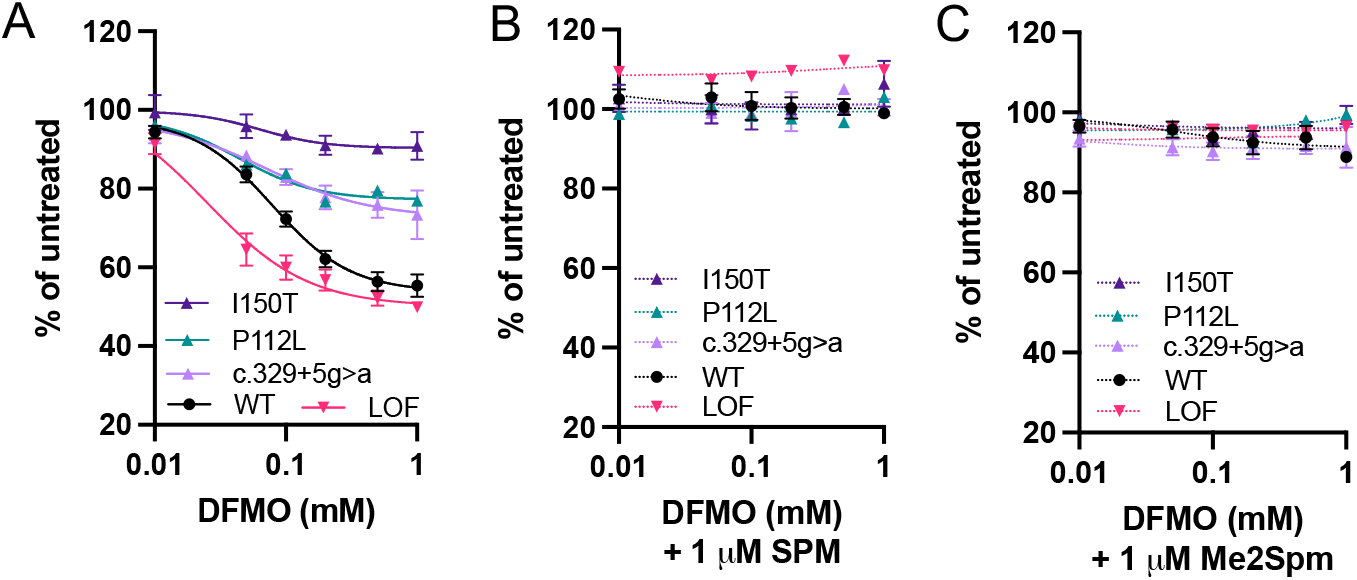
Effects of DFMO on SRS patient fibroblasts: mutation-specific sensitivity and rescue by spermine. Curves depict DFMO-mediated growth inhibition of fibroblast cell lines derived from healthy donors (WT) or SRS patients, as indicated by *SMS* mutation. Cells were grown for 96 h in the presence of increasing doses of DFMO alone (**A**) or in combination with low doses of SPM (**B**), or 1,12-Me_2_SPM (**C**). “LOF” indicates a mutation causing complete loss of SMS enzymatic activity. Data points indicate the means with SE of triplicate determinations of at least 2 biological replicates.

**Table 1.** Differential sensitivity of SMS variant fibroblast cell lines to DFMO (t = 96 h).

| *SMS genotype:* | *WT* | *I150T* | *P112L* | *c.329+5* | *LOF* |
| --- | --- | --- | --- | --- | --- |
| IC<sub>50</sub> (μM) | 79.1 | 63.7 | 53.2 | 71.2 | 27.3 |
| % max inhibition | 46.2 | 9.7 | 24.0 | 27.4 | 48.8 |

Regardless of the degree of growth inhibition resulting from DFMO exposure, proliferation was completely rescued when low doses of SPM or its mimetic (1,12-Me_2_SPM) were included during treatment (**Fig. 3B-D**). These results indicate that the most common SRS-associated mutations, all of which are hypomorphic, increase resistance to the growth inhibitory effects of DFMO. Additionally, as extracellular sources of SPM include the diet and microbiota, growth inhibition by DFMO is not anticipated to be a treatment limitation in SRS.

### DFMO induces uptake of and cooperates with extracellular spermine or a mimetic to reduce the SPD/SPM ratio

We previously established that both lymphoblastoid and fibroblast cell lines originating from SRS patients could import SPM from the extracellular environment to restore a more normal polyamine distribution profile (18,19). Inhibition of polyamine biosynthesis with inhibitors such as DFMO is known to evoke a compensatory increase in polyamine import activity. We therefore investigated the effects of treating SRS cells with DFMO in the presence of physiologically relevant concentrations of extracellular SPM. Aminoguanidine (1 mM) was included in the culture medium to inhibit the extracellular oxidation of SPM (26). In SRS lymphoblasts with the *SMS*^*G56S*^ mutation, providing exogenous SPM in a concentration range relevant to that reported in human plasma (27) did not reduce the concentration of SPD or total polyamines, but it did reduce the SPD/SPM ratio (**Fig. 4A**). However, in the presence of DFMO, these same SPM concentrations more effectively restored intracellular SPM concentrations, decreased SPD levels, and returned the SPD/SPM ratio to that observed in SRS WT cells.

**Fig. 4.**
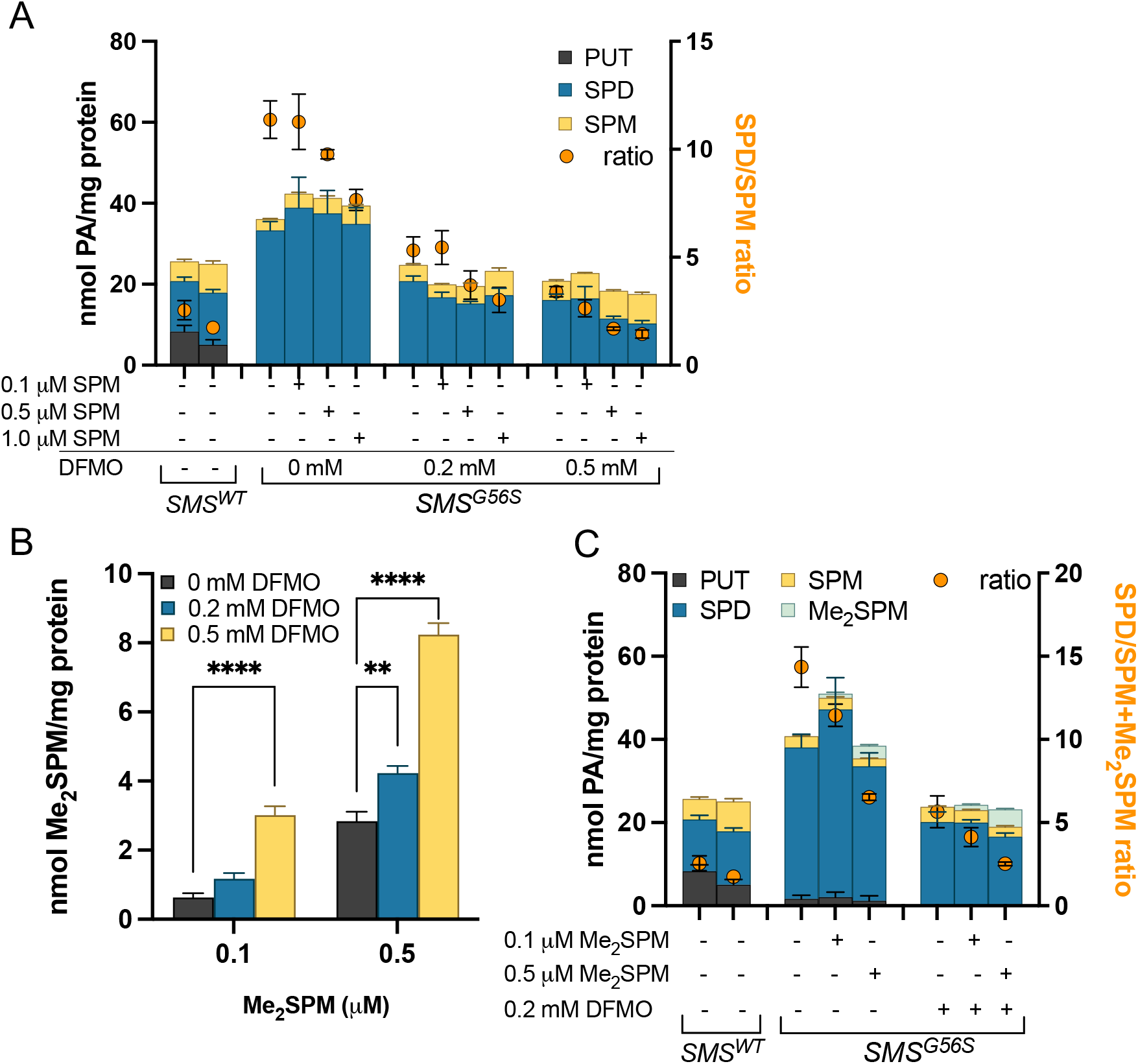
SPD/SPM ratio reduction by DFMO is enhanced in the presence of extracellular spermine or a spermine mimetic. (**A)** *SMS*^*G56S*^ lymphoblasts were exposed to physiologically relevant doses of extracellular spermine in the presence of DFMO (and 1 mM aminoguanidine) for 96 h. (**B)** DFMO treatment increases cellular uptake of the SPM mimetic Me_2_SPM. (**C)** Low-dose DFMO improves response to Me_2_SPM, reducing SPD, improving uptake, and returning the ratio (orange dots plotted on the right axis) to that of *SMS*^*WT*^ cells. In **A** and **C**, intracellular polyamine levels are plotted on the left axis and indicate the means with SE of duplicate determinations of at least 2 biological experiments.

To clarify the effect of DFMO treatment on polyamine uptake, we replaced SPM with 1,12-Me_2_SPM, a SPM mimetic that is resistant to amine oxidase activity and is capable of supporting cellular proliferation in the presence of DFMO (28-30). Me_2_SPM is also easily distinguished from endogenous SPM by HPLC, thus allowing precise measurement of intracellular concentrations. The presence of DFMO increased the accumulation of intracellular Me_2_SPM by approximately 3-fold (**Fig. 4B**), thereby reducing the dose required and potentially reducing off-target effects. Polyamine pool analyses confirmed that DFMO and Me_2_SPM cooperatively reduced the elevated levels of SPD, total polyamines, and the SPD/SPM ratio in SRS cells, indicating the combination of DFMO and Me_2_SPM as a potential treatment option.

### DFMO extends lifespan in a Drosophila model of SRS

We used a previously reported *Drosophila* model of SRS to determine the potential for phenotype reversal by DFMO *in vivo (31)*. The phenotype of this model includes a significantly shortened lifespan that can be reversed by overexpression of wild-type human *SMS*. As the *dSMS* gene is autosomal in flies, both male and female flies were included in the study. The addition of increasing concentrations of DFMO to the feed of *dSms*^*-/-*^ flies increased lifespan in a dose-dependent manner (**Figs. 5, S3**). Survival of male flies was significantly extended starting at 1 mM DFMO, with the highest dose, 10 mM, lengthening median life span from 21 days to 35 days (**Fig. 5B**). Survival in female flies was also significantly extended, albeit to a lesser extent (**Fig. S3**). These results indicate the feasibility of oral administration of DFMO in eliciting systemic, phenotype-reversing effects at the organismal level, and together with our biochemical data in patient cell lines, support the use of DFMO clinically in the context of SRS.

**Fig. 5.**
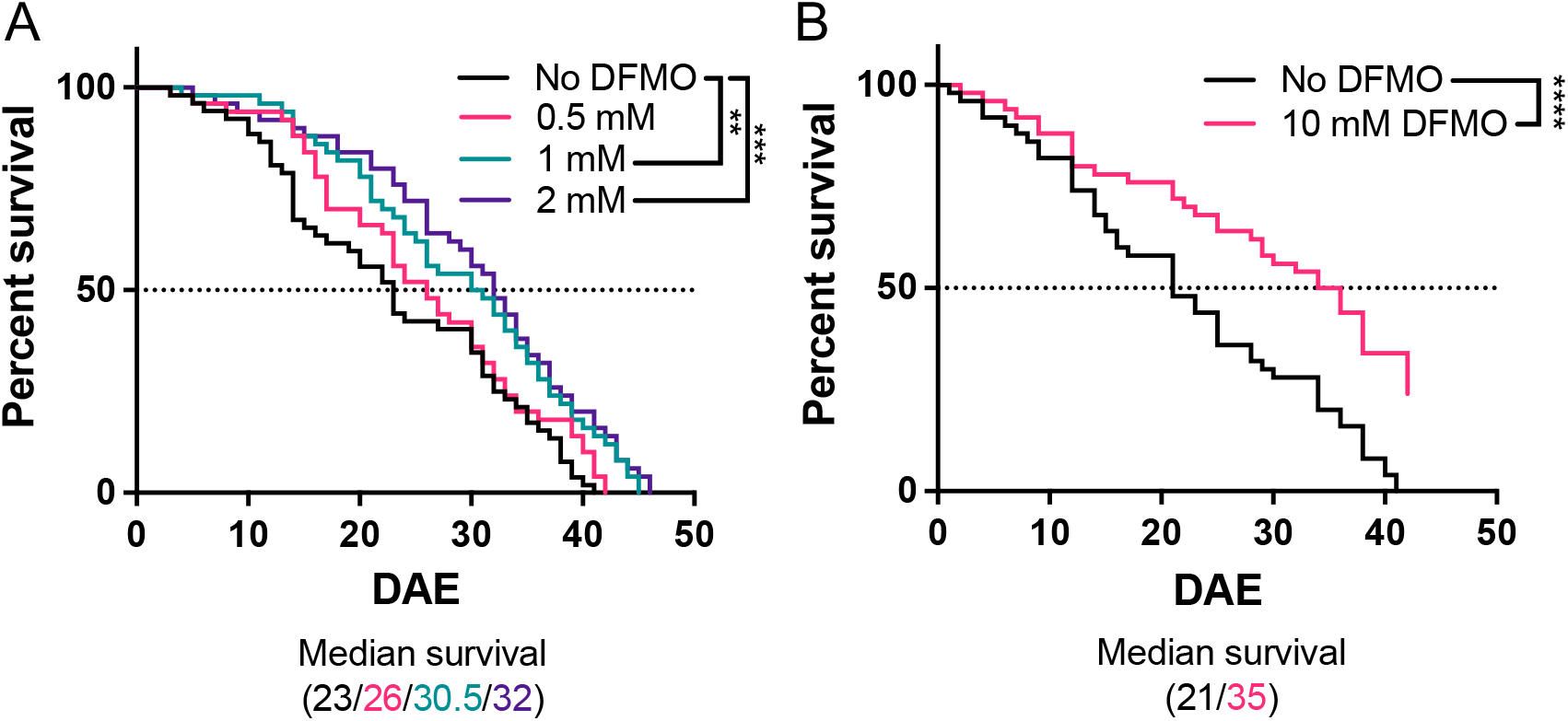
DFMO extends lifespan in male *dSms*^*-/-*^ *Drosophila*. The indicated concentration of DFMO was administered in the feed and survival was observed. In (A), n = 52 in the control group, and 50 in each of the 0.5, 1, and 2 mM DFMO treatment groups; in (B), n = 50 each in the 0 and 10 mM DFMO treatment groups. The resulting survival curves were compared using the log-rank (Mantel-Cox) test. DAE, days after eclosure.

## Discussion

Our studies indicate the potential repurposing of DFMO as a treatment strategy for males with SRS. To date, only a single patient has been identified with complete loss of SMS activity (14). Partial loss-of-function (LOF), hypomorphic variants are much more common, which often interfere with the dimerization required for efficient aminopropyltransferase activity (32-34). Our treatment of SRS cells in culture with DFMO demonstrates that the hypomorphic SMS enzymes can indeed convert SPD into SPM, a reaction that depends upon induction of AMD1. In contrast, treatment of the SMS LOF line reduces the SPD concentration but fails to produce SPM. Increased availability of dcSAM to the hypomorphic enzyme likely contributes to this reaction; however, whether the SMS enzyme itself is affected by the DFMO treatment remains to be determined. It should be noted that although DFMO is a specific and irreversible inhibitor of ODC, the synthesis and rapid turnover of ODC results in some PUT generation which, in the presence of high dcSAM is rapidly converted to SPD and ultimately SPM, if functional SMS is present (35).

Decarboxylated SAM, resulting from AMD1 activity, provides the aminopropyl donor for transfer onto PUT or SPD acceptor molecules, thereby forming SPD or SPM, respectively. The increase in basal AMD1 enzyme activity observed in the SRS lymphoblasts compared to those with wild-type SMS is consistent with the increased generation of dcSAM. Of the polyamines, SPM most potently down-regulates AMD1 activity (36,37), suggesting that the reduced SPM levels characteristic of SRS cells may contribute to these results. The reduced use of dcSAM by the variant SMS enzymes further contributes to its accumulation. Similar results have been reported in cells from SRS patient families including those with the *SMS*^*G56S*^ variant and the *SMS*^*I150T*^ variant, not included in the current study (24). In Gy mice, which completely lack the *Sms* gene, significantly elevated dcSAM levels were detected in the brain, kidney, liver, and heart (24).

Though compensatory responses to polyamine biosynthesis inhibition by DFMO has been well documented in a variety of cell types (36-38), it has not been previously investigated in SRS patient cells. DFMO treatment greatly induced AMD1 activity in lymphoblastoid cell lines. While the extent of induction was similar between WT cells and the G56S variant, induction in the c.329+5 variant significantly exceeded either of these. However, the resulting accumulation of the AMD1 product, dcSAM, did not correlate with AMD1 activity, but rather inversely correlated with SMS activity, as indicated by the resulting positive correlation with SPD/SPM ratio (note that SPD is entirely depleted by DFMO in WT cells, thus no ratio is indicated in **Fig. 2A**). By inhibiting ODC, DFMO reduces the production of PUT and SPD, thereby reducing the pool of potential acceptors of dcSAM in aminopropyltransferase reactions. This substrate limitation, combined with active SPD and SPM synthesis, accounts for the lower dcSAM accumulation in WT cells. Though the elevated SPD levels in SRS cells reduces the substrate limitation, the lack of efficient aminopropyltransferase activity by the SMS variants allows additional dcSAM buildup that is available for transfer mainly onto SPD by SMS. The c.329+5 g>a *SMS* variant disrupts a splice site, resulting in a truncated protein in addition to the full-length, functional SMS protein due to read-through (5). Consequently, cells with this variant have measurable SMS activity that maintains a SPD/SPM ratio approximately double that of WT controls (our study and (5)). DFMO facilitates the conversion of SPD into SPM at concentrations as low as 50 μM in both lymphoblasts and fibroblasts with this variant. This increased utilization of dcSAM in the SMS reaction is likely responsible for the difference in dcSAM measured between the 2 SRS cell lines. The G56S variant of SMS possesses very little enzymatic activity, resulting in a ratio approximately 6-fold greater than controls (our study and (10)) and requiring higher DFMO concentrations to appropriately reduce the ratio, although it should be noted that the 0.2 mM concentration of DFMO used remains much lower than the doses commonly used to inhibit biosynthesis in cultured cells. The evident inverse association between intracellular SPM and dcSAM concentrations following DFMO treatment may also reflect negative regulation of AMD1 activity by SPM *in situ* (37).

Though dcSAM levels are elevated in SRS patient cells, no evident differences were detected in baseline SAM levels, as previously observed (24). However, following treatment with DFMO, a clear decrease in SAM was noted in the *SMS*^*G56S*^ cell line, concurrent with this cell line demonstrating the greatest accumulation of dcSAM. This result warrants further consideration and potentially indicates a need for SAM supplementation in the clinical context, as SAM is an important methyl group donor for many reactions.

Our studies indicate the potential for DFMO to improve the low bioavailability of SPM that is apparent with its oral ingestion in SMS-deficient animals (15,16). Additionally, while DFMO cannot stimulate the production of SPM by SMS LOF variants, it does stimulate uptake of SPM, as indicated by our rescue experiments, thereby providing a source of SPM to lower the SPD/SPM ratio. DFMO also improved the uptake and response to *R,R*-1,12-Me_2_SPM, a metabolically stable SPM mimetic we previously reported to improve the SPD/SPM ratio in SRS patient cell lines (19). Alpha-methylation of SPM prevents its acetylation and/or oxidation via the polyamine catabolic enzymes, cellular monoamine and diamine oxidases, as well as serum amine oxidases (30,39). The production of toxic byproducts via these reactions limits the use of native SPM as a treatment *in vivo*, especially when administered to animals by intravenous or intraperitoneal (i.p.) routes (40-42). Thus, as Me_2_SPM is amenable to more direct modes of delivery, it could serve as an alternative to replace SPM. Although Me_2_SPM effectively reduced SPD/SPM ratios when administered to intraperitoneally mice, evidence of toxicity was observed at higher concentrations (19). Our data indicating a combined effect of DFMO and Me_2_SPM importantly imply that, when used together, lower doses of each will be required to elicit the desired effect, thereby reducing the potential for adverse effects.

Importantly, DFMO is clinically approved for the treatment of African trypanosomiasis (43), and its use as a maintenance therapy for pediatric neuroblastoma patients has provided valuable data on safety and administration (44,45). DFMO has a decades-long history of safe administration in clinical trials and has recently been repurposed to treat patients with Bachmann-Bupp syndrome (BABS), a newly identified genetic disorder resulting from an ODC gain-of-function variant (46,47). The presentation of BABS emulates that of SRS in several aspects, including developmental delay, hypotonia, dysmorphic features, and non-specific brain MRI findings (48), most of which dramatically improved with DFMO treatment (46). Our studies in SRS patient-derived cell lines indicate that DFMO effectively reverses the aberrantly elevated SPD/SPM ratio that is the main biochemical hallmark of SRS. Additionally, adding DFMO to the feed of *Drosophila* lacking *dSms* significantly extends lifespan, a main phenotype in this SRS model (31). Taken together, our results indicate that repurposing DFMO for the treatment of SRS is a valid therapeutic strategy that targets the underlying biochemistry driving the SRS phenotype, thereby supporting further clinical translation of DFMO into this population.

### Experimental procedures

#### Cell lines, culture conditions, and compounds

Lymphoblastoid and dermal fibroblast cell lines were derived from male SRS patients or healthy donors at the Greenwood Genetic Center (Greenwood, SC) and were previously described (5,10,12,18). Lymphoblastoid lines were maintained in RPMI 1640 containing 15% fetal bovine serum (Gemini Bio-Products, Sacramento, CA), 2 mM glutamine, sodium pyruvate, non-essential amino acids (NEAA), and penicillin/streptomycin at 5% CO_2_ and 37ºC. Fibroblast cell lines were maintained in DMEM containing 15% fetal bovine serum, L-glutamine, sodium pyruvate, NEAA, and antibiotics at 5% CO_2_ and 37ºC. (*R,R*)-1,12-Me_2_SPM was synthesized previously (49) and DFMO was provided by Professor Patrick M. Woster (Medical University of South Carolina).

#### Polyamine concentration determinations and enzyme activity assays

Intracellular polyamine concentrations of cell or tissue lysates were determined by HPLC following acid extraction and dansylation of the supernatant, as previously described (50). HPLC standards included diaminoheptane (internal standard), PUT, SPD, SPM, and acetylated SPD and SPM, all of which were purchased from Sigma Chemical Co. Enzyme activity assays were performed for SAMDC using radiolabeled SAM, as previously described (36). Enzyme activities and intracellular polyamine determinations are presented relative to total cellular protein, as determined using the Bio-Rad Protein Assay (Hercules, CA) with interpolation on a bovine serum albumin standard curve.

#### SAM and dcSAM quantification

SAM and dcSAM levels were quantified using a method developed by the Analytical Pharmacology Shared Resource of the Sidney Kimmel Comprehensive Cancer Center at Johns Hopkins. Decarboxylated SAM-d3 was purchased from Toronto Research Chemicals (North York, Ontario, CA). Unlabeled dcSAM and SAM were purchased from Sigma Chemical Co. Deionized water was obtained from a Millipore Milli-Q-UF filtration system (Milford, MA). A standard solution containing the target analytes, SAM and dcSAM (1 mg/mL), was prepared in acetonitrile:water (50:50, v/v). These stock standard solutions were diluted with methanol:water (50:50, v/v) to prepare a mixed working solution with different concentrations. dcSAM-d3 was used as internal standard (I.S.) and prepared in water at a concentration of 15 ng/mL. Stock and working solutions were stored at 4°C for daily use.

Methanol:water (50:50, v/v) was selected as the extraction solvent and yielded good extraction recoveries for SAM and dc-SAM. First, 500 μl of the extraction solvent were added to the frozen cell pellets. The samples were slowly thawed at room temperature. Subsequently, the cells were shock-frozen in -80°C for 10 min and thawed for a second time at room temperature. The samples were vortexed and sonicated for 60 sec, then vortexed and centrifuged at 12000 rpm for 5 min at 4°C. After centrifugation, 50 μL of each supernatant was transferred into an autosampler vial along with 20 μL of the internal standard for LCMS/MS analysis of dc-SAM. Another 50 μL of the supernatant was added to 450 μL of cold methanol, and then 50 μL of the diluted solution along with 20 μL of the internal standard was transferred into autosampler vials for LCMS/MS analysis of SAM.

Separation was achieved with a Waters Acquity UPLC BEH Amide 2.1 × 100 mm, 1.7 μm (Waters Corporation, Milford, MA). Mobile phase A was 0.1% formic acid in 10 mM of ammonium formate, and mobile phase B was 0.1% formic acid in acetonitrile. A gradient was used and mobile phase B was held at 15% with a flow rate of 0.15 mL/min for 0.5 minutes. It was then increased to 80% over 1.5 minutes and held there for 2 minutes, then returned to 15% B and was held there for one minute. The column effluent was monitored using an SCIEX 6500 triple quadrupole mass-spectrometric detector (Sciex, Foster City, CA, USA) using electrospray ionization operating in positive mode. The mass spectrometer was programmed to monitor the following MRM transitions: 399.04 → 250.10 for SAM, 356.96 → 250.20 for dcSAM, and 357.97 → 250.00 for the internal standard, dcSAM-d3. Calibration curves for SAM and dcSAM were computed using the area ratio peak of the analysis to the internal standard by using a quadratic equation with a 1/x^2^ weighting function and calibration ranges of 5 to 1000 ng/mL. For SAM, the additional 1:10 dilution was factored into the final concentration calculation.

#### Cell proliferation assays

Fibroblast cells were seeded at 1000 cells/well in 96-well plates and allowed to attach overnight. Growth medium was replaced with that containing increasing concentrations of DFMO with or without 1 μM *R,R*-1,12-Me_2_SPM or SPM in the presence of 1 mM aminoguanidine (100 μL total per well), and cells were incubated 96 h. CellTiter-Blue reagent (Promega, Madison, WI) was added (20 μL/well) followed by an additional 3-h incubation at 37ºC. Fluorescence was measured in white, opaque-bottom plates at 560_Ex_/590_Em_ on a SpectraMax M5 platereader (Molecular Dynamics, Sunnyvale, CA), using wells containing medium alone (no cells) as background controls. Results are presented as percentages relative to untreated cells. Individual points represent the means with standard error of at least 2 independent biological experiments, each performed in triplicate wells.

#### Drosophila studies

The *dSms* mutant fly (d*Sms*^*-/-*^) strain has been previously characterized (31,51). Flies were maintained in vials with cornmeal-molasses-yeast medium at 22°C, 65% humidity, and 12-hours light / 12-hours dark. To measure the life span, newly eclosed flies were collected, and 10-20 flies of the same sex were kept in vials with food with or without the indicated concentration of DFMO. Flies were transferred to new vials every week, and the number of dead flies was counted every other day.

#### Statistical analyses

All statistically significant differences were determined using GraphPad Prism software (v. 9.3.1, La Jolla, CA). Dose responses of fibroblast cell lines to DFMO were plotted by non-linear regression for determination of IC_50_ values. Differences were determined using mixed-effects analysis with correction for multiple comparisons using the Bonferroni method. Comparisons between untreated and treated WT and SRS cell lines were performed by 2-way ANOVA with correction for multiple comparisons by the Tukey method. \**p* values ≤ 0.05 were considered statistically significant; \*\**p* ≤ 0.01, \*\*\**p* ≤ 0.001, **** *p* ≤ 0.0001.

## FOOTNOTES

These studies were supported by funding from the Million Dollar Bike Ride, Orphan Disease Center at the University of Pennsylvania (MDBR-20-135-SRS and MDBR-21-106-SRS to R.A.C. and T.M.S.), the Chan-Zuckerberg Initiative, the National Institutes of Health National Cancer Institute (R01CA204345, R01CA235963 to R.A.C.), and the Eunice Kennedy Shriver National Institute of Child Health and Human Development (R01HD110500 to R.A.C. and T.M.S.). Synthesis of 1,12-Me_2_SPM was supported by the Russian Science Foundation (Grant 17-74-20049 to A.R.K.). The project was also supported by the Analytical Pharmacology Shared Resource of the Sidney Kimmel Comprehensive Cancer Center at Johns Hopkins directed by Dr. Michelle Rudek [NIH grants P30CA006973 and UL1TR003098, and the Shared Instrument Grant S10RR026824] and grant number UL1TR003098 from the National Center for Advancing Translational Sciences (NCATS), a component of the National Institutes of Health (NIH), and the NIH Roadmap for Medical Research. Its contents are solely the responsibility of the authors and do not necessarily represent the official view of the NCATS or NIH.

