## Supplemental Figures 1 - 3 for "Difluoromethylornithine rebalances aberrant polyamine ratios in Snyder-Robinson syndrome: mechanism of action and therapeutic potential"

### Slide 1
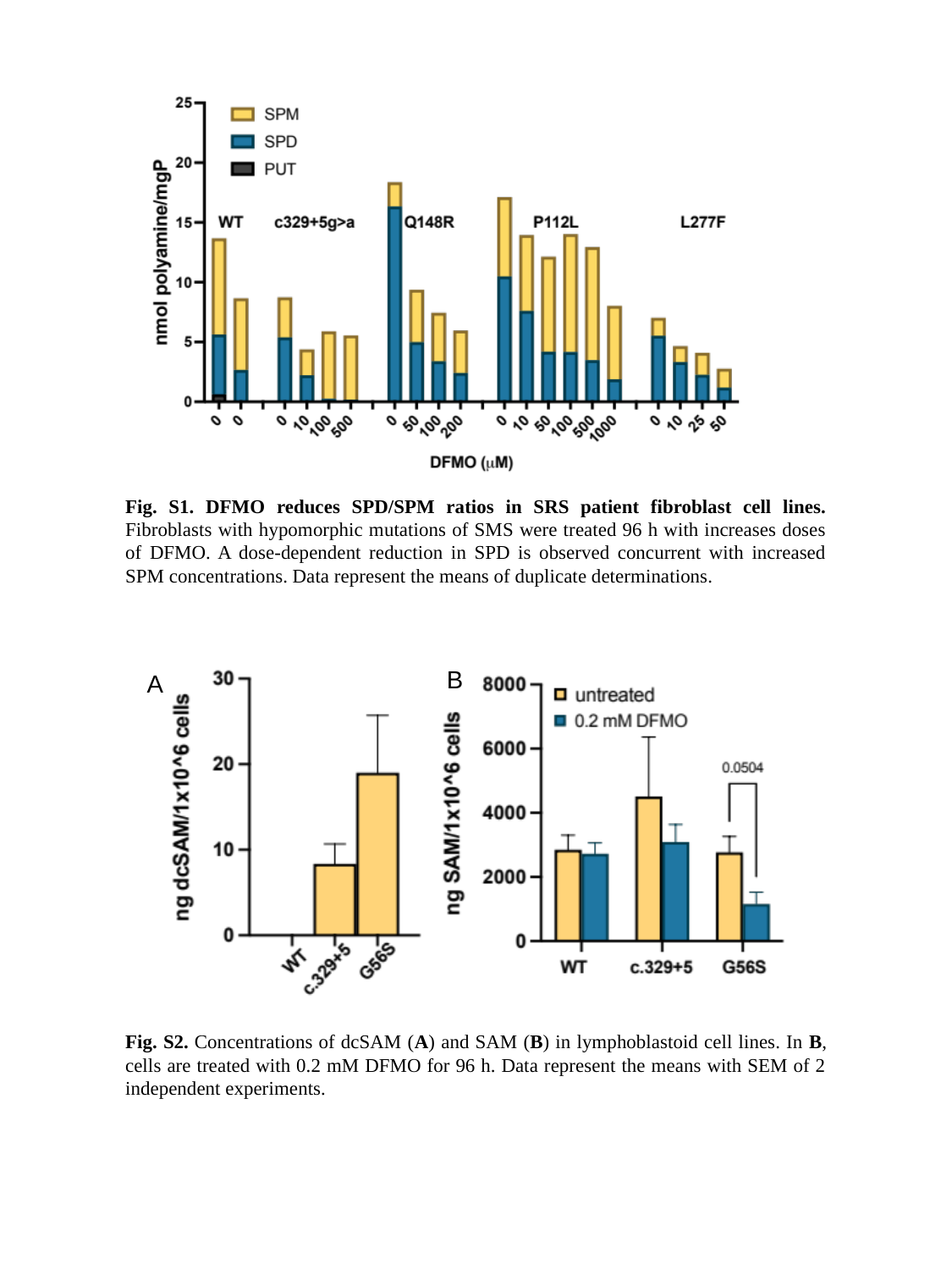

Fig. S1. DFMO reduces SPD/SPM ratios in SRS patient fibroblast cell lines. Fibroblasts with hypomorphic mutations of SMS were treated 96 h with increases doses of DFMO. A dose-dependent reduction in SPD is observed concurrent with increased SPM concentrations. Data represent the means of duplicate determinations.
B
A
Fig. S2. Concentrations of dcSAM (A) and SAM (B) in lymphoblastoid cell lines. In B, cells are treated with 0.2 mM DFMO for 96 h. Data represent the means with SEM of 2 independent experiments.

### Slide 2
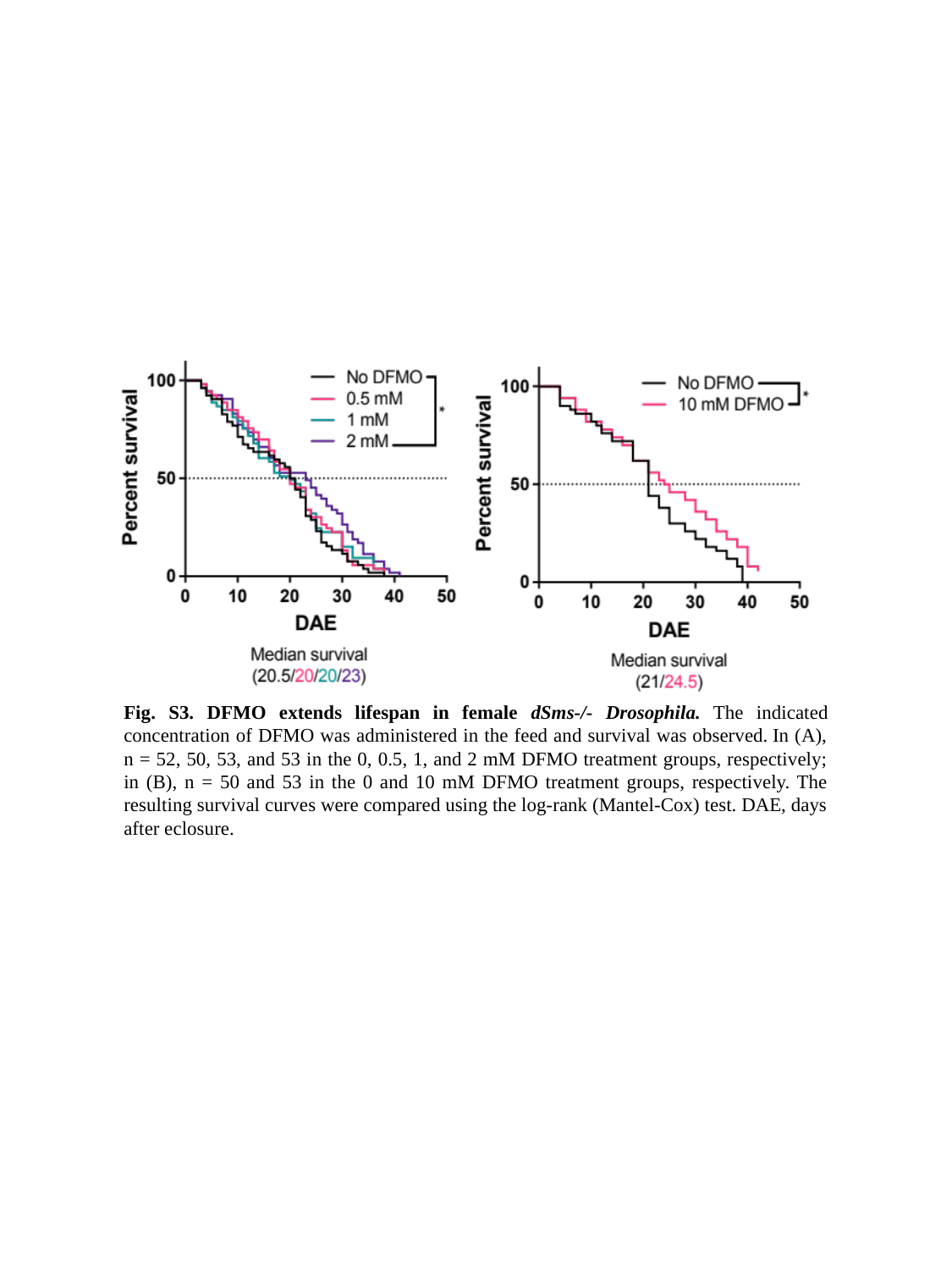

B
A
Fig. S3. DFMO extends lifespan in female dSms-/- Drosophila. The indicated concentration of DFMO was administered in the feed and survival was observed. In (A), n = 52, 50, 53, and 53 in the 0, 0.5, 1, and 2 mM DFMO treatment groups, respectively; in (B), n = 50 and 53 in the 0 and 10 mM DFMO treatment groups, respectively. The resulting survival curves were compared using the log-rank (Mantel-Cox) test. DAE, days after eclosure.
